# Hematological phenotypes in GATA2 deficiency syndrome arise from secondary injuries and maladaptation to proliferation

**DOI:** 10.1101/2024.09.24.614663

**Authors:** Juncal Fernandez-Orth, Cansu Koyunlar, Julia M. Weiss, Emanuele Gioacchino, Hans de Looper, Geoffroy Andrieux, Mariëtte Ter Borg, Joke Zink, Irene Gonzalez-Menendez, Remco Hoogenboezem, Baris Yigit, Kirsten J Gussinklo, Roger Mulet-Lazaro, Charlotte Wantzen, Sophie Pfeiffer, Christian Molnar, Eric Bindels, Sheila Bohler, Mathijs Sanders, Leticia Quintanilla-Martinez, Marcin Wlodarski, Melanie Boerries, Ivo P. Touw, Charlotte Niemeyer, Miriam Erlacher, Emma de Pater

## Abstract

The GATA2 transcription factor is a pivotal regulator of hematopoiesis. Disruptions in the *GATA2* gene drive severe hematologic abnormalities and are associated with an increased risk of myelodysplastic syndromes and acute myeloid leukemia; however, the mechanisms underlying the pathophysiology of GATA2 deficiency remain still unclear. We developed two different mouse models that are based on serial and limiting donor cell transplantation of (aged) GATA2 haploinsufficient cells and mirror the symptoms of GATA2 deficiency. Similar to what has been observed in patients, our models show that GATA2 haploinsufficiency leads to B lymphopenia, monocytopenia, lethal bone marrow failure (BMF), myelodysplasia and leukemia. Leukemia arises exclusively as a result of BMF, driven by somatic aberrations and accompanied by increased *Myc* target expression and genomic instability. These findings were confirmed in human GATA2+/− K562 cell lines showing defects in cytokinesis and are in line with the fact that monosomy 7 and trisomy 8 are frequent events in patients with MDS.

**Key points:**

- In a mouse model for GATA2 deficiency, leukemia emerges from bone marrow failure
- Maladaptation to proliferative signals and chromosomal segregation defects contribute to the hematological phenotypes in GATA2 deficiency

## Introduction

GATA2 plays an essential role in hematopoietic stem and progenitor cell (HSPC) development and differentiation^1^. Patients with *GATA2* germline mutations suffer from monocytopenia, B and NK cell lymphopenia, neutropenia and an inversion of the CD4/CD8 T cell ratio but most notably, bone marrow failure (BMF)^2,3^ and are predisposed to myelodysplastic syndrome (MDS) and acute myeloid leukemia (AML)^4^. Although several genetic alterations are known to cause BMF and predispose to myeloid neoplasia^5,6^, GATA2 deficiency syndrome stands out because of its high risk of malignant transformation, with 80% of patients aged 40 having developed MDS/AML^7^. So far, more than 400 *GATA2* germline mutations have been identified^1,8–10^, but a clear genotype-phenotype correlation could not be established due to high patient variability^7,11–13^.

GATA2 is expressed in immature stem and progenitor cells, as well as in mature hematopoietic cell types such as megakaryocytes, mast cells and monocytes^14–17^, and is essential for the generation and maintenance of hematopoietic stem cells (HSCs) in the aorto-gonad-mesonephros region in the murine embryo^18–22^.

In heterozygous mouse embryos and adults, HSPCs defined as lineage marker negative, Sca^+^Kit^+^ (LSK) cells are significantly reduced^23^ in number and functionality^24,25^. Interestingly, several phenotypes found in GATA2 deficiency can be recapitulated in zebrafish, where the *GATA2* gene is duplicated into *Gata2a* and *Gata2b*. Whereas Gata2a is mostly expressed in endothelia and required for generation of the first HSPCs through endothelial-to-hematopoietic transition (EHT), Gata2b is expressed mostly in HSPCs after EHT. Complete loss of Gata2b is non-lethal in zebrafish and results in neutropenia^26,27^. Heterozygous mutations in *Gata2b* result in dysplasia in adult zebrafish^28,29^. Interestingly, the conserved +9.5 enhancer of GATA2, which is also mutated in some patients and required for embryonic and adult hematopoiesis^30^, drives *Gata2a* mRNA expression. A complete deletion of this enhancer results in monocytopenia, and occasionally, acute leukemia^31^.

Here, we characterized two distinct mouse models and a human cell model to investigate GATA2 haploinsufficiency. Our models recapitulate GATA2-deficiency associated hematological phenotypes such as B lymphopenia and monocytopenia and provide a unique opportunity to study leukemogenesis in this disease setting. In our model, leukemia emerges exclusively as a result of severe BMF, accompanied by increased *Myc* target expression and genomic instability.

## Methods

### Mouse maintenance

*Gata2^+/−^* mice^19^ were maintained on a C57BL/6 background and genotyped as previously described^24^. *Vav-cre;Gata2^fl/+^* mice^32^ were kindly provided by Elaine Dzierzak (Edinburgh, Scotland) and genotyped as described. B6.SJL-Ptprca Pepcb/BoyJ mice served as recipients. Animals were housed under specific-pathogen-free conditions. All experiments were performed in accordance with animal welfare regulations. **(Primers, Suppl. Table 1)**

### BM sampling

Mouse bone marrow (BM) was obtained by flushing femurs, tibias and pelvis using a 27G (BD) needle with PBS supplemented with 5 IU/mL penicillin (P), 5 μg/mL streptomycin (S), and 10% fetal calf serum (PS-FCS) and stored on ice until use.

### LSK cells isolation for transplantation or *in vitro* experiments

Filtered BM cell suspension was depleted of lineage-marker positive cells (#130-090-858, Miltenyi Biotec) according to manufactureŕs instructions. Lineage negative cells were stained with Sca1+cKit+ antibodies plus a viability dye to exclude dead cells (eFluor™ 506), and sorted accordingly (FACSAria Fusion, FACSAria III and MoFlo Astros EQ, BD Bioscience). For *in vitro* experiments, LSK cells were cultured in 96-well U-bottom plates (25,000 cells/well in 200 μL IMDM medium (Sigma Aldrich) in PS-FCS and recombinant murine cytokines stem cell factor (SCF), thrombopoietin (TPO) and FLT3L (100 ng/ml each, ImmunoTools, Friesoythe) at 5% CO2 and 37°C.

### Transplantation experiments

For Figure 2, 3×10^6^ freshly isolated nucleated BM CD45.2^+^ cells from 65-week-old *Gata2^+/−^* mice were transplanted via tail vein injection into lethally irradiated (10.5Gy) CD45.1^+^ recipient mice. Donor cell chimerism was determined in PB. 4 months after transplantation cells were retransplanted into new lethally irradiated recipients.

**Figure 1:**
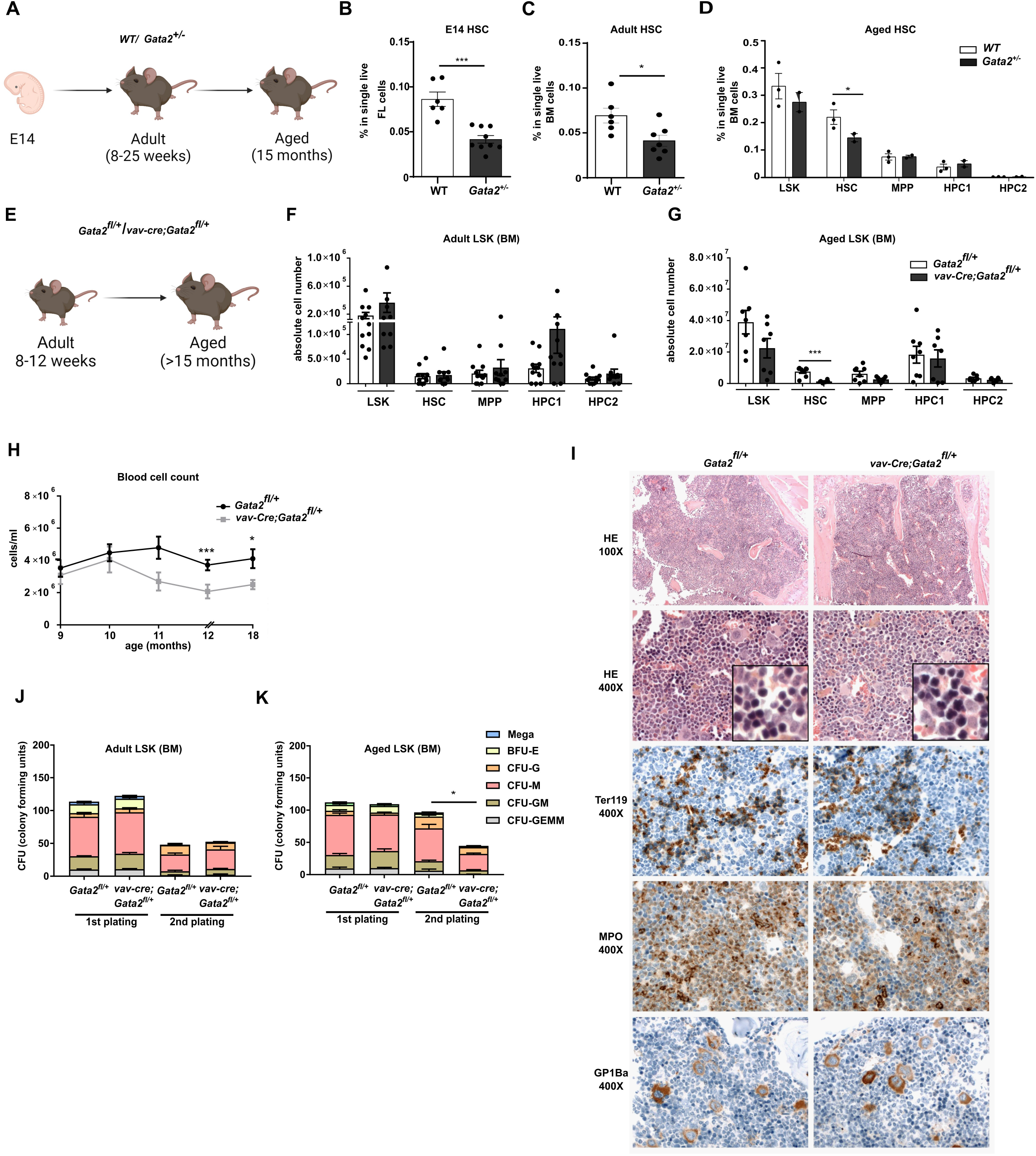
Aged Gata2^+/−^ and vav-cre;Gata2^fl/+^ mice exhibit diminished hematopoietic stem cell functionality together with significant defects when subjected to proliferative stress. (A) Representative example of the developmental stage and genotype of the mice used in the study. % of hematopoietic stem cells (HSC) in both WT (white) and Gata2^+/−^ (grey) from embryo (E14) (B), adult (8-25 weeks) (C) and aged (15 months) (D) mice. Representative example of the developmental stage and genotype of the mice used in the study (E). Number of stem and progenitor cells in adult (F) *Gata2^fl/+^* (white) and *vav-Cre;Gata2^fl/+^* (grey) and aged mice (G). Blood cell count of *Gata2^fl/+^* and *vav-Cre;Gata2^fl/+^* over the time (H). Histopathology performed on sternum from aged *Gata2^fl/+^*and *vav-Cre;Gata2^fl/+^* mice (I). Adult HSC cells from both *Gata2^fl/+^* and *vav-Cre;Gata2^fl/+^* adult mice were serially cultured in colony forming assays (CFU) (J, first plate; K, replating). (GMEM: granulocyte, monocyte, erythrocyte, megakaryocyte, GM: granulocyte, macrophage progenitor, M: monocytic precursor cells, G: granulocyte precursor cell, BFU-E: Burst-forming unit-erythroid).

**Figure 2:**
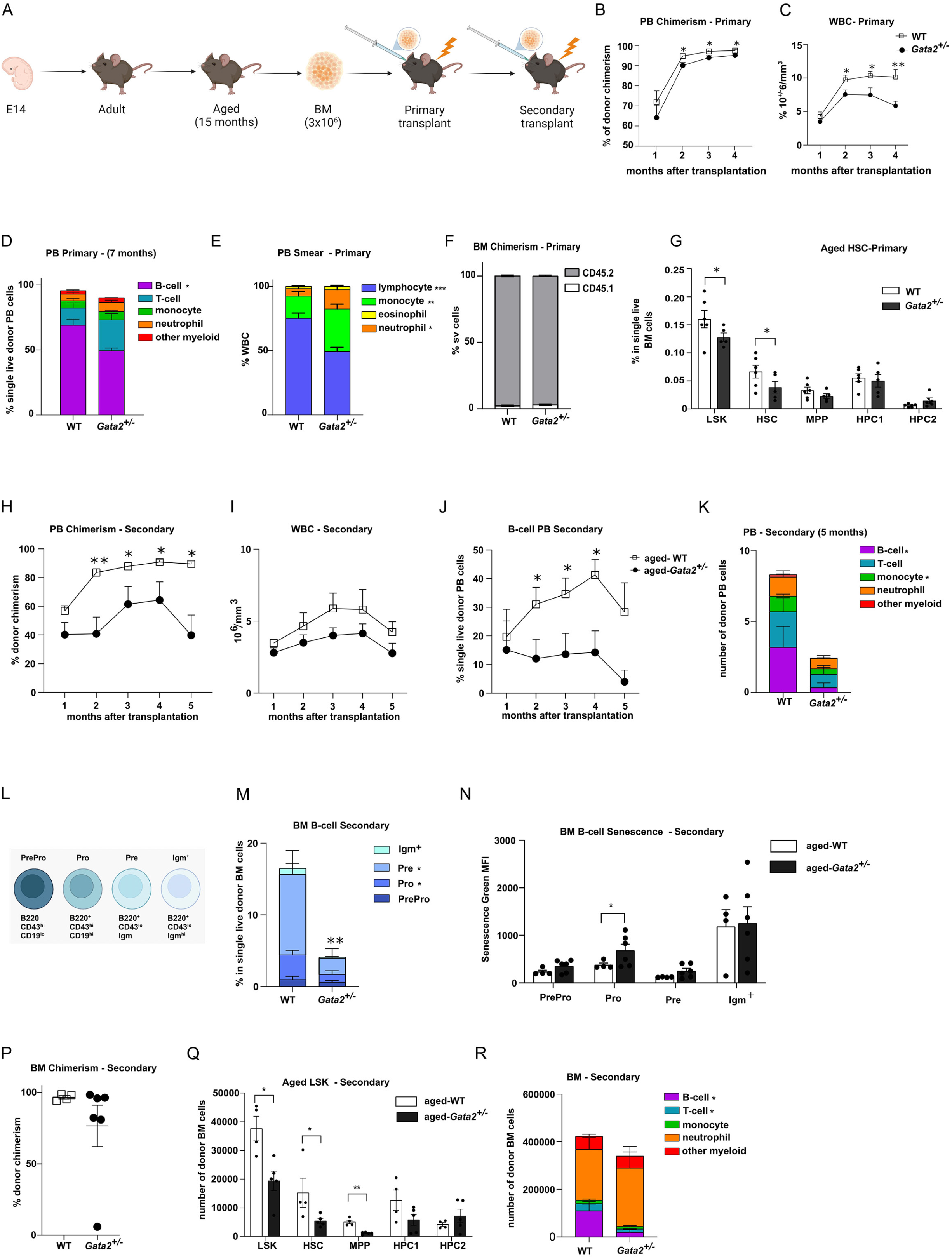
Characterization of aged *Gata2^+/−^* transplanted mice. **(A)** Representative example of the transplantation assay performed with aged WT and *Gata2^+/−^* mice. (**B)** Peripheral blood chimerism analysis performed over the months in WT and *Gata2^+/−^* mice. (**C)** Total white blood cell count analyzed over the time in WT and Gata2^+/−^ mice. (**D)** Analysis of the different blood cell subsets performed in PB 7 months after primary transplantation in WT and *Gata2^+/−^* mice. (**E)** Analysis of the different blood cells subsets from PB smears 7 months after primary transplantation in WT and *Gata2^+/−^* mice. **(F)** BM chimerism analyzed 7 months after transplantation. (**G)** Analysis of the aged HSC BM compartment 7 months after primary transplantation in WT and *Gata2^+/−^* mice. (**H)** Representative example of the secondary transplantation assay performed with aged WT and *Gata2^+/−^* mice. **(I)** Peripheral blood chimerism analysis performed in WT and *Gata2^+/−^* mice. **(J)** Total white blood cell count analyzed over the time in WT and *Gata2^+/−^*mice. **(K)** B cell analysis from PB 4 months after secondary transplantation in WT and *Gata2^+/−^* mice. (**L)** Analysis of the different blood cell subsets performed in PB 4 months after secondary transplantation in WT and *Gata2^+/−^* mice. **(M)** Schematic representation of B cell differentiation in WT and *Gata2^+/−^* mice. **(N)** B cell differentiation in BM after secondary transplantation in WT and *Gata2^+/−^* mice. **(O)** B cell senescence analyzed 4 months after secondary transplantation in WT and *Gata2^+/−^* mice. **(P)** BM chimerism analyzed after secondary transplantation in WT and *Gata2^+/−^* mice. **(Q)** Analysis of aged HSC BM compartment 4 months after secondary transplantation. **(R)** Analysis of the different blood cell subsets performed in PB 4 months after secondary transplantation in WT and *Gata2^+/−^* mice.

Similarly, 25,000 LSK cells isolated from adult (8-12 weeks old) *Gata2^fl/+^* or *vav-cre;Gata2^fl/+^* mice were transplanted into CD45.1+ recipients. Six weeks after first transplantation, 25,000 CD45.2+ LSK cells were re-transplanted into secondary recipients.

For Figure 3, CD45.1+ mice recipients were lethally irradiated (9.5 Gy) and reconstituted by retro-orbital injection of 20,000 LSK cells isolated from 8-12 weeks Gata2^fl/+^ or vav-cre;Gata2^fl/+^ mice.

**Figure 3:**
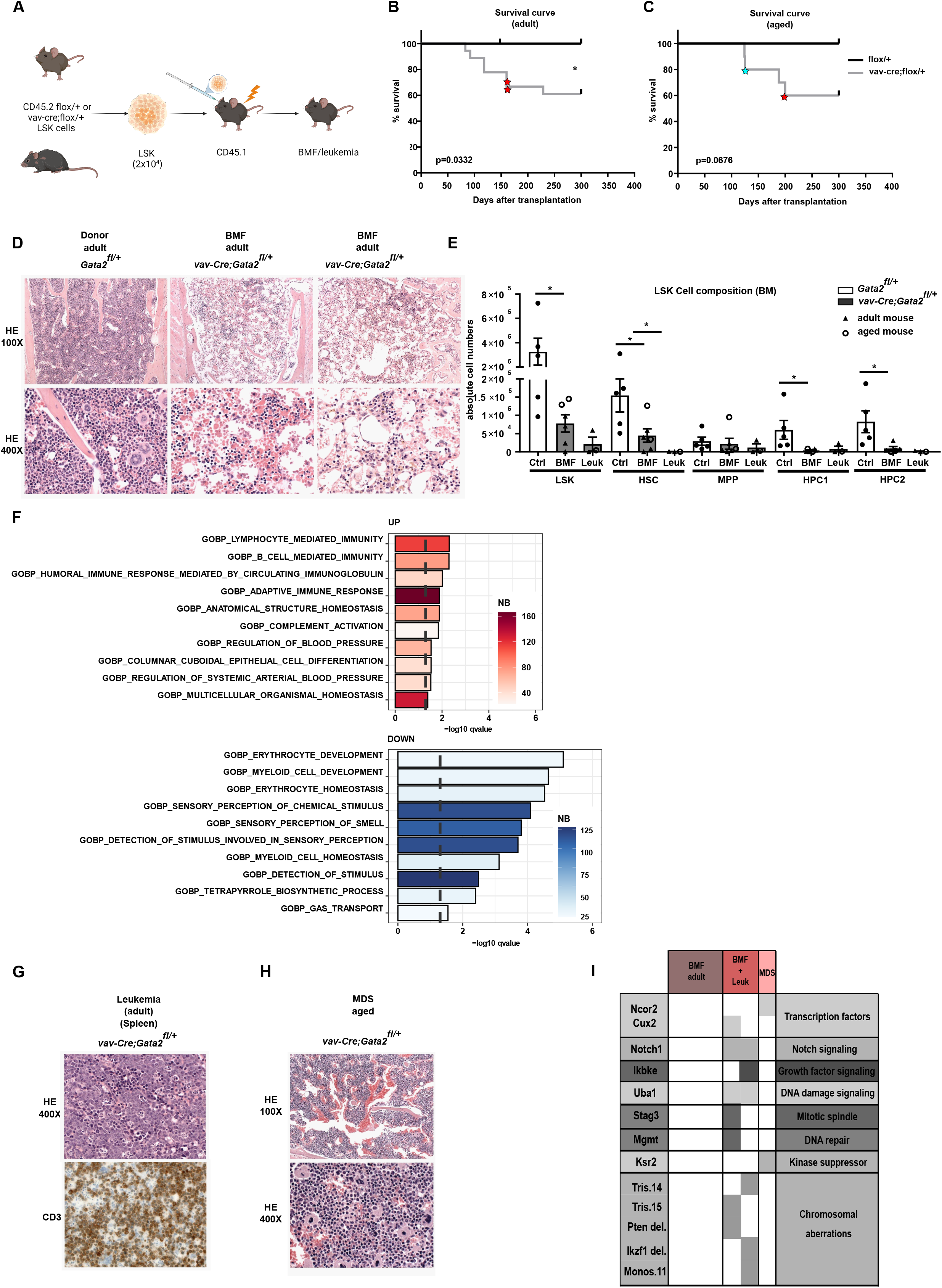
Transplantation of *vav-cre;Gata2^fl/+^* LSK into lethally irradiated mice leads to BMF and leukemia. **(A)** Representative example of the transplantation assay performed with *Gata2^fl/+^* and *vav-Cre;Gata2^fl/+^* mice. Survival curves obtained after subjecting adult **(B)** and aged **(C)** *Gata2^fl/+^* and *vav-Cre;Gata2^fl/+^* mice to transplantation and followed over the time. Red stars show the mice that succumbed due to leukemia. **(D)** H&E stainings performed on sternum obtained from donor and succumbed mice due to BMF leukemic and myelodysplasic mice at 100X and 400X. (**E)** Analysis of the different BM cell subsets from animals that succumbed due to BMF and leukemia compared to *Gata2^fl/+^* mice. (**F**) GSEA of RNAseq analysis of CD45.2+ BM cells from *Gata2^fl/+^* and *vav-Cre;Gata2^fl/+^* mice. X-axis indicates significance value. (**G)** H&E and CD3 IHC performed on spleen obtained from an adult succumbed mouse due to leukemia. **(H)** H&E stainings performed on BM obtained from an aged succumbed mouse due to MDS. **(I)** WES and array-CGH analysis performed on BM samples obtained from succumbed animals due to BMF, leukemia and MDS.

### Cell staining

All hematopoietic organs were incubated for 15 min in red blood cell lysis buffer (150 mM NH4Cl, 10 mM NaHCO3, 1 mM EDTA) and stained with antibodies described in **Suppl. table 2**. For intracellular antigens, cells were fixed and permeabilized with Cytofix/Cytoperm Kit (BD Biosciences).

Cell proliferation was determined using CellTrace™ CFSE Cell Proliferation Kit, (Thermo Fisher Scientific) and analyzed via Flow Cytometry after 72 or 96 hours of culture at 37°C with 5% CO2 in IMDM, P/S-FCS together with (Flt3L, TPO and SCF) or without cytokines. Apoptosis was determined either by using FITC Annexin V Apoptosis Detection Kit I (BD Biosciences) according to manufacturer instructions or by combining Annexin-V with 4,6-diamidino-2-phenylindole (DAPI) (Molecular Probes). Cell cycle analysis was performed on fixed cells stained with 4,6-diamino-2-phenylindole (DAPI) and ki56 FITC (Thermofisher) 1:25. Cells were acquired via Flow Cytometry (LSR Fortessa, FACS Aria 3, LSR 2, BD Bioscience) and analyzed with FlowJo (FlowJo LLC, Becton Dickinson). The gating strategy is shown in **(Suppl. Fig 1)**.

### Colony-forming unit assay

500 freshly sorted cells were seeded in MethoCult GF M3434 (Stem Cell Technologies) and colonies were scored after 10 days. Afterwards, MethoCult was dissolved using PBS and up to 10^4^ cells were re-plated.

### Histopathology

Murine sterna were fixed in 4% formalin, decalcified in EDTA and paraffin embedded. Sections (3-5 µm) were stained with haematoxylin and eosin (HE). Immunohistochemistry was performed on an automated immunostainer (Ventana Medical Systems) according to the company’s protocols. Antibodies are listed in **Suppl. Table 2**. Pictures were taken on an Axioskop 2 plus Zeiss microscope equipped with a Jenoptik and ProgRes C10.

### RNA Sequencing and gene set enrichment analysis (GSEA)

RNA was extracted with Trizol from BM-derived lineage marker negative cells with the RNeasy Micro kit (Qiagen). Sequencing libraries were prepared using the TruSeq Stranded mRNA Library Prep Kit (Illumina) or SMARTer Ultra Low RNA kit (Clontech) and sequenced on both the hiseq and Novaseq 6000 instruments (Illumina) and analysed as described previously^31–36^.

### Statistics

Data are presented as mean ± SEM. All statistical analysis was carried out in GraphPad Prism 8.0.1 (GraphPad Software Inc.). Normally distributed data were analyzed using the non-parametric Mann-Whitney test. A *p* value less than 0.05 was considered significant.

## Results

### Ageing in Gata2 heterozygous mice is associated with loss and functional defects of HSCs

In individuals with monoallelic germline *GATA2* mutations, the risk of hematological complications rises with age^7^. Therefore, we studied the hematopoietic system of *Gata2^+/−^* mice from embryonic stages to 15 months of age. The *Gata2^+/−^* HSPC compartment **(Figure 1A)** had fewer phenotypic HSCs in the embryonic fetal liver and BM of adult (8-25 weeks) and aged mice (15 months) compared to wildtype (WT) mice **(Figure 1B-D)**. No other hematological phenotypes such as cytopenias were detected in either adult or aged *Gata2^+/−^* mice **(not shown)**.

To investigate whether the BM HSC reduction was caused by hematopoietic-specific functions of Gata2, we deleted one *Gata2* allele selectively in the hematopoietic compartment (*vav-cre;Gata2^fl/+^*), leading to GATA2 haploinsufficiency after HSC generation^37^ **(Figure 1E)**. At the age of 8-12 weeks, adult *vav-cre;Gata2^fl/+^* mice exhibited neither a decrease in BM HSC numbers **(Figure 1F)** nor any other discernible hematological phenotype compared to control *Gata2^fl/+^* mice **(Suppl. Figure 2A)**. However, aged *vav-cre;Gata2^fl/+^* mice (>15 months) showed a significant loss of HSCs **(Figure 1G),** accompanied by mild cytopenia, which was initially observed at 11-12 months of age **(Figure 1H).** No other alteration of the BM compartment of 60 weeks old *vav-cre;Gata2^fl/+^*mice was observed compared to *Gata2^fl/+^* mice **(Suppl. Figure 2B).** Histology of the sternum showed mild dysplasia of the erythroid lineage **(Figure 1I)**. This suggests that the decrease in HSCs observed in embryonic and adult *Gata2^+/−^* mice harboring a germline mutation is primarily caused by defects during embryonic HSC development and/or microenvironmental influences. In contrast, HSC loss during ageing is an intrinsic cellular effect of GATA2 haploinsufficiency.

To functionally characterize GATA2 haploinsufficient HSPCs, we performed colony forming assays with serial replating. Adult *Gata2^+/−^* HSCs, defined as CD48^-^CD150^+^LSK cells, did not show any differentiation defect (**Suppl. Figure 2C-D**). Similarly, replating of adult *vav-cre;Gata2^fl/+^* LSK cells did not reveal any differentiation defect **(Figure 1J).** In contrast, significantly fewer *vav-cre;Gata2^fl/+^*than *Gata2^fl/+^* colonies were formed after replating of aged LSK cells **(Figure 1K).** The lack of erythroid, monocytic and granulocytic cells (GEMM) *vav-cre;Gata2^fl/+^* colonies in secondary plates indicated a compromised self-renewal ability of aged *vav-cre;Gata2^fl/+^* stem cells.

### B lymphopenia and monocytopenia after transplantation of aged Gata2^+/−^ bone marrow

Despite the functional defects of aged HSCs, no pathologies such as BMF or leukemia were seen in aged *Gata2^+/−^* or *vav-cre;Gata2^fl/+^*mice. This might be explained by the lack of proliferative stress, caused by environmental stimuli. Therefore, we challenged the aged hematopoietic system by serial transplantations of 3×10^6^ BM cells isolated from 15 months *Gata2^+/−^* and WT donor mice (**Figure 2A**). Primary BM transplantation of aged-*Gata2^+/−^* cells resulted in a mild but significant reduction of donor chimerism in peripheral blood (PB) **(Figure 2B).** Total white blood cell (WBC) count was significantly decreased **(Figure 2C)**, mostly due to a decrease in the B cell compartment compared to WT transplanted mice **(Figure 2D-E)**, recapitulating the most reproducible phenotype of GATA2 deficiency patients^38^. Total BM chimerism was not altered, indicating a selective differentiation defect of the B cell lineage **(Figure 2F)**. Notably, the HSC compartment was significantly reduced in mice transplanted with aged *Gata2^+/−^* BM compared to aged WT transplanted mice **(Figure 2G)**. After secondary transplantation **(Figure 2A)**, donor chimerism in PB was consistently reduced **(Figure 2H)**, although recipient cells now compensated WBC numbers **(Figure 2I)**. Donor-derived B lymphopenia persisted in secondary recipients transplanted with aged-*Gata2^+/−^*BM **(Figure 2J)**. In addition, monocytopenia was now observed in aged *Gata2^+/−^* transplanted mice when assessing donor-derived cells **(Figure 2K)**.

B cell differentiation was blocked due to increased senescence in pro-B cells in aged *Gata2^+/−^* BM transplanted mice compared to WT **(Figure 2L-N)**. This block was only partial and, even though IgM^+^ class switched B-cells were reduced, they remained detectable as previously seen in patients^39^. BM cellularity was comparable in both groups 5 months after secondary transplantation **(Suppl. figure 3B)**, but BM chimerism was reduced in 3 out of 6 mice transplanted with aged *Gata2^+/−^* BM, indicating HSC exhaustion **(Figure 2P)**. Supporting this finding, a significant reduction in the absolute numbers of MPPs and HSCs in BM of mice after secondary transplantation with aged-*Gata2^+/−^* BM was found **(Figure 2Q)**. T cell differentiation was significantly reduced in BM of mice transplanted with aged-*Gata2^+/−^* BM compared to WT **(Figure 2R).** The number and differentiation of myeloid and erythroid cells in the BM of aged *Gata2^+/−^*BM secondary transplanted mice were not altered despite the monocytopenia found in PB **(Figure 2R, Suppl. Figure 3A)**.

To show that the phenotypic changes are a result of the aged *Gata2^+/−^* hematopoietic system, similar experiments were performed using LSK cells isolated from 8 to 16 week old *vav-cre;Gata2^fl/+^* mice **(Suppl. Figure 3C)**. Marginal differences in organ cell counts of spleen and LN were observed in primary and secondary recipients of *vav-cre;Gata2^fl/+^* LSK cells **(Suppl. Figure 3D-E),** while no other phenotypes were identified **(Suppl. Figure 3F and not shown)** confirming that the aging of *Gata2^+/−^* HSPCs is important for the functional defects, recapitulating cytopenias frequently observed in patients.

### Gata2 haploinsufficiency predisposes to lethal BMF and leukemia

In a second approach, we challenged the haploinsufficient GATA2 hematopoietic system by transplanting low numbers (i.e. 2×10^4^) of sorted LSK cells into lethally irradiated WT recipients. Donor LSK cells were derived from both adult (8-12 weeks) and aged (54 weeks) *Gata2^fl/+^* and *vav-cre;Gata2^fl/+^* mice **(Figure 3A)**. Recipient mice were monitored closely for up to one year. In both groups, adult and aged *Gata2^fl/+^*and *vav-cre;Gata2^fl/+^* donor cells engrafted successfully (**adult**: *Gata2^fl/+^* n=10; *vav-cre;Gata2^fl/+^*, n=18; **aged**: *Gata2^fl/+^* n=7; *vav-cre;Gata2^fl/+^* n=10). However, 10 recipients transplanted with *vav-Cre;Gata2^fl/+^* LSK cells had to be sacrificed between 100 and 200 days after transplantation because of a severe deterioration of their condition (**adult**: n=6 of 18, **aged**: n=4 of 10). Median time from transplantation to death was 118 days (range 83 to 160 days) in adult and 160 days (range 125 to 200) in aged mice. In contrast, all mice from both ages transplanted with *Gata2^fl/+^*LSK cells remained healthy **(Figure 3B-C).** Analysis of succumbed *vav-Cre;Gata2^fl/+^* recipients revealed BMF in 6 animals (**adult**: n=4, **aged**: n=2). Histopathological and flow cytometric analysis of their sternum showed aplastic BM **(Figure 3D)** and significant alterations in the HSPC compartment **(Figure 3E)** respectively, together with a reduction in almost all hematopoietic lineages **(Suppl. Figure 4A)**. RNA-seq of Ly5.2^+^ BM cells isolated 14 days post transplantation revealed a significantly reduced differentiation signature of both erythroid and myeloid lineages in *vav-Cre;Gata2^fl/+^* cells compared to WT **(Figure 3F)**. This suggests that the lineage differentiation defects are cell intrinsic and are driven by defects occurring quickly after transplantation.

Three BMF mice (**adult**: n=2, **aged**: n=1) presented with donor-derived T cell leukemia **(red stars in Figure 3B-C)** with thymic and splenic involvement **(spleen, Figure 3G)**. All three had few leukemic cells in the otherwise empty BM (**Suppl. Figure 4A**). One of the succumbed mice transplanted with aged *vav-Cre;Gata2^fl/+^*cells showed a normocellular dysplastic megakaryo– and erythropoiesis, left shifted myelopoiesis and increased presence of B– and T-cells **(Figure 3H)**. All remaining recipients were sacrificed one year after transplantation. Similar to the characteristics observed in non-transplanted aged donors, there was a discernible loss of HSPCs, MPPs and HSCs **(Suppl. Figure 4B)** without other alterations in the BM cell composition **(Suppl. Figure 4C)**. On the other hand, mice transplanted with both adult and aged *Gata2^fl/+^*HSPCs cells did not show any hematological pathologies. HSPCs derived from one donor were transplanted into several recipients, using multiple donors (**adult n=5 donors, 18 recipients, aged: n=5 donors, 10 recipients**). Importantly, HSPCs from the same donor resulted in diverse pathologies in recipient mice, irrespective of the donor’s age, suggesting that leukemic events arose after transplantation rather than before **(Suppl. Figure 4D).**

To investigate the mechanisms of leukemic transformation and MDS development, WES and array-CGH were performed (**adult** (BMF n=3, BMF+leukemia n=1) and **aged** (BMF+leukemia n=1, MDS n=1). While no mutations were identified in donor mice (n=2) or animals exhibiting BMF, the somatic events were detected in mice with leukemia or MDS. In addition to mutations in oncogenes and tumor suppressors, chromosomal aberrations were detected in all analyzed mice presenting with leukemia, irrespective of whether they were transplanted with adult or aged *vav-Cre;Gata2^fl/+^* LSK cells (**Figure 3H**). We identified cases with trisomy 15 (encompassing the *Myc* locus and corresponding to human trisomy 8) and a *Pten* deletion. Additionally, both cases (2 out of 2) with leukemia had *Notch1* mutations, which are linked to T-ALL leukemia in humans^40^, along with *Uba1* mutations found also in humans with MDS and autoinflammatory symptoms^41^.

Of note, clonal outgrowth of cells with genetic alterations resulting in acute T cell leukemia occurred exclusively after BMF in our model. Whether the normocellular MDS found in one recipient was preceded by BMF is unclear.

### Delayed hematopoietic reconstitution of Gata2^+/−^ LSK cells precedes bone marrow failure

Our mouse models suggested that proliferative stress contributes to GATA2 deficient phenotypic onset. To better understand the kinetics of hematopoietic failure, we analyzed reconstitution at different time points following transplantation **(Figure 4A)**. Homing of HSPCs to their niche, as determined 15 hours after transplantation, was not affected by GATA2 haploinsufficiency **(Figure 4B)**. However, between day 4 and 14 after transplantation, significantly fewer *vav-cre;Gata2^fl/+^*HSPCs were found in the recipient mice compared to *Gata2^fl/+^* **(Figure 4C),** together with a significant reduction in the HSC fraction **(Figure 4D)** as well as a trend in the different mature cells **(Figure 4E).** *Vav-cre;Gata2^fl/+^* HSPCs underwent apoptosis significantly more than *Gata2^fl/+^* cells 14 days post transplantation shown by the increased percentage of sub-G_0_ apoptotic cells **(Figure 4F).** Although no significant differences were found in the cell cycle stages of LSK cells, few cells were found in G_0_ stage, indicating the high proliferative pressure on the LSK pool of *Gata2^fl/+^* cells **(Figure 4G)** at this time point. The colony forming capacity of *vav-cre;Gata2^fl/+^* LSK cells isolated 14 days post transplantation was significantly impaired **(Figure 4H)**. Surprisingly, 42 days post transplantation, LSK cells were found at normal or even increased numbers in all recipient mice **(Figure 4I).** This compensatory LSK expansion between week 2 and 6 after transplantation was corroborated by a decrease in the number of apoptotic cells **(Suppl. Figure 4E)** without differences in the mitotic cell stages **(Suppl. Figure 4F)**. In summary, recipients of *vav-Cre;Gata2^fl/+^* cells successfully reconstituted their hematopoietic system before developing BMF, although reconstitution was significantly delayed. It is conceivable that the initial loss of LSK cells, followed by a period of increased proliferation, resulted in clonal hematopoiesis paving the way for hematopoietic exhaustion at later time points.

**Figure 4:**
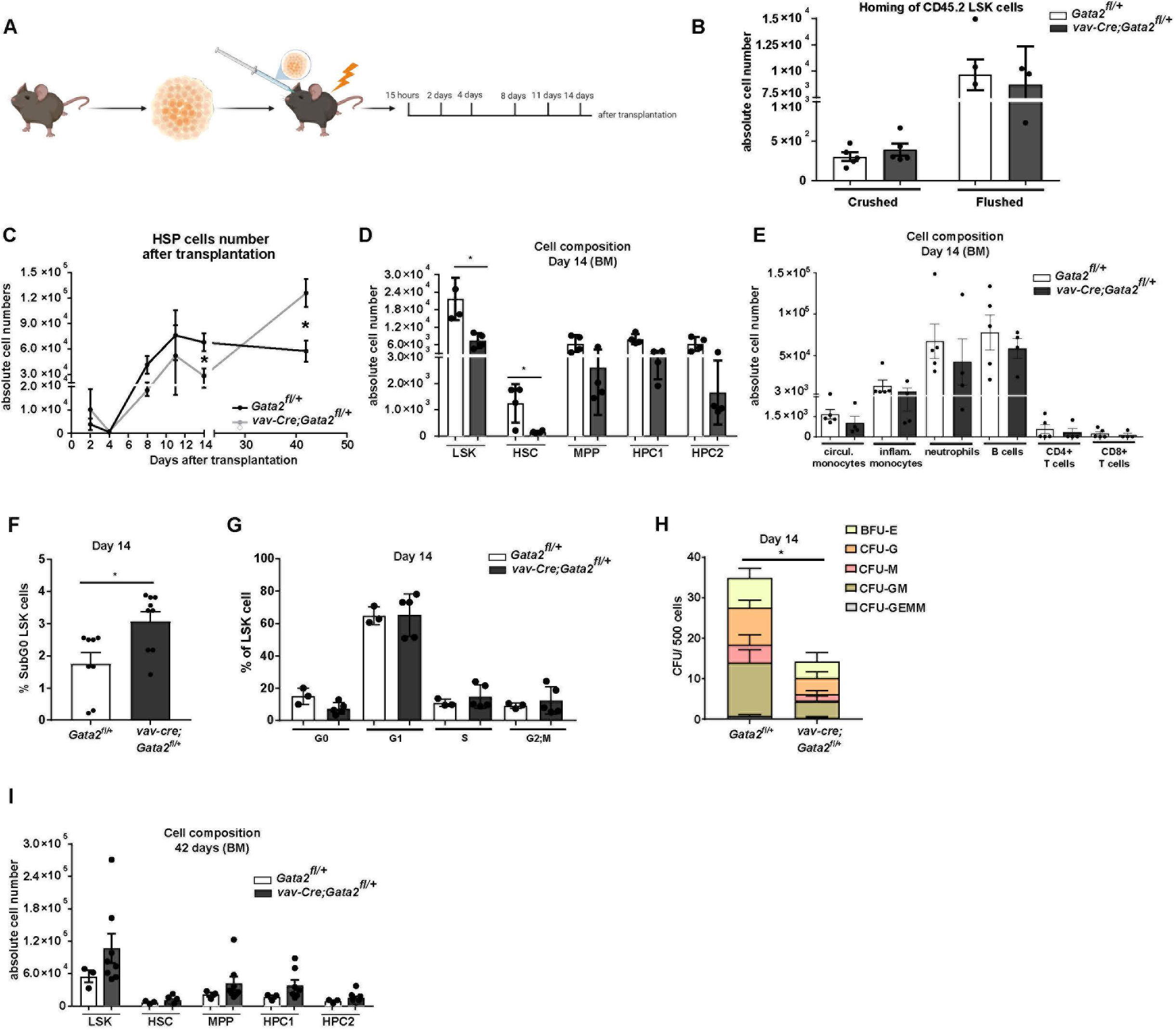
*vav-cre;Gata2^fl/+^* cells lead to a delayed hematopoietic reconstitution. **(A)** Representative example of the time points analyzed after transplantation with *Gata2^fl/+^* and *vav-Cre;Gata2^fl/+^* mice. **(B)** Absolute cell numbers obtained 15 hours after transplantation after crushing or flushing the bones. **(C)** Absolute LSK numbers analyzed over the time after transplantation. **(D)** Analysis of the different BM cell subsets from *Gata2^fl/+^* and *vav-Cre;Gata2^fl/+^* mice 14 days after transplantation. **(E)** Analysis of the different blood cell subsets 14 days after transplantation in *Gata2^fl/+^*and *vav-Cre;Gata2^fl/+^* mice. **(F)** % of SubG0 LSK cells after Ki67/DAPI staining performed in *Gata2^fl/+^* and *vav-Cre;Gata2^fl/+^* mice. **(G)** % of mitotic cell stages performed in *Gata2^fl/+^* and *vav-Cre;Gata2^fl/+^* mice. **(H)** Colony forming assays from *Gata2^fl/+^* and *vav-Cre;Gata2^fl/+^* LSK cells 14 days after transplantation. **(I)** Analysis of the HSC BM compartment from both *Gata2^fl/+^* and *vav-Cre;Gata2^fl/+^* mice 42 days after transplantation.

### GATA2-associated phenotypes are accompanied by increased Myc target gene expression, proliferation defects and genomic instability

To dissect the molecular consequences of GATA2 deficiency, transcriptomes of WT and *Gata2^+/−^* HSCs at various developmental stages were investigated. The most prominent gene set that was enriched in *Gata2^+/−^* HSCs at E14, adult and aged HSCs was *Myc Targets V1* (**Figure 5A-G**). GSEA analysis was visualized using Cytoscape^42^. This revealed that native and primary transplanted aged HSCs also upregulate gene sets related to nucleotide excision repair and DNA damage in *Gata2^+/−^*compared to WT **(Suppl. Figure 5A-B)**. Related to the immune deficiencies seen in these mice, we observed a deregulation of the B cell regulatory pathway (**Suppl. Figure 5B**) as well as in both myeloid and lymphoid pathways in the HSPC1 compartment **(not** shown).

**Figure 5:**
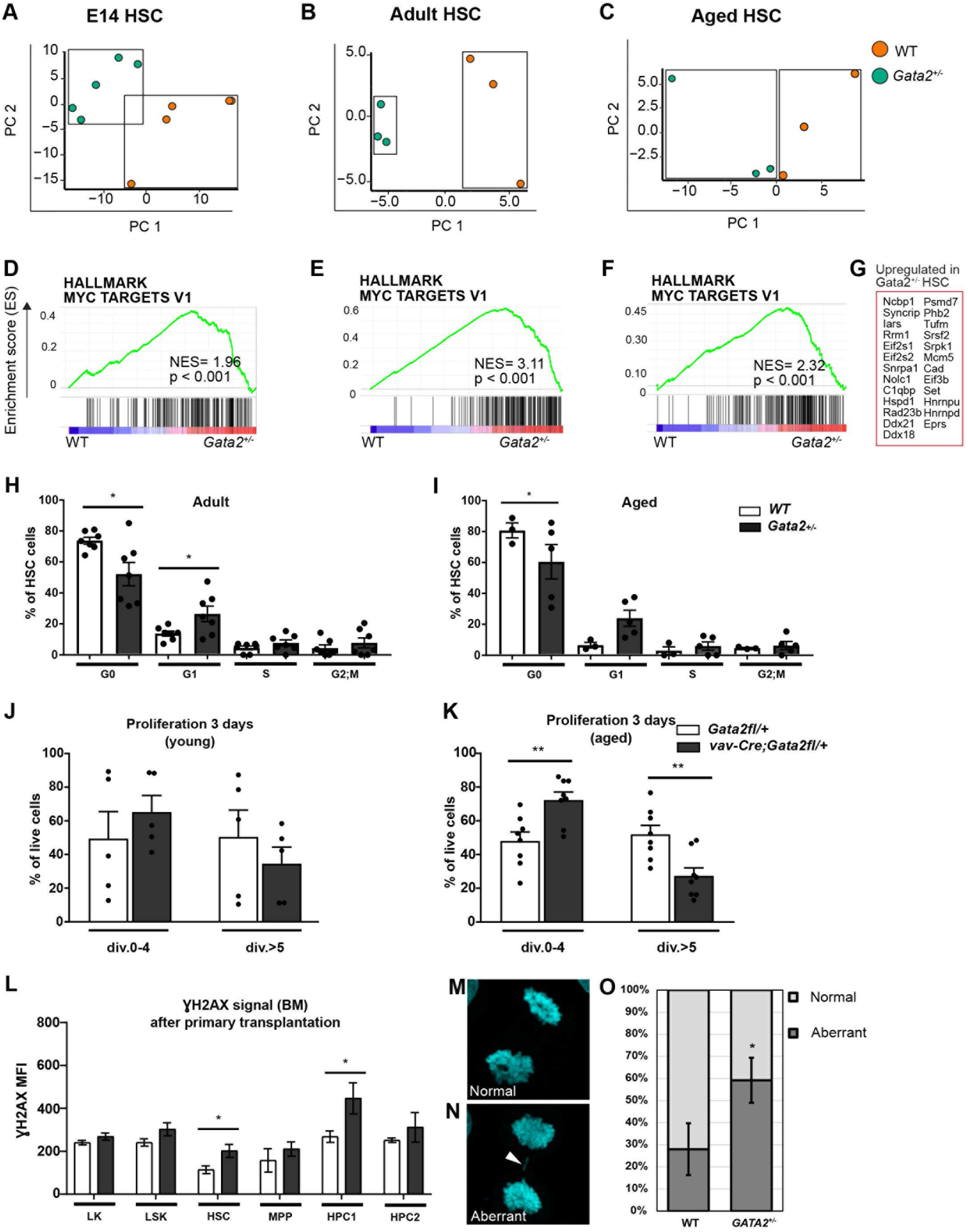
*Gata2* associated phenotypes are accompanied by proliferation defects and genomic instability. **(A)** PCA of E14 **(B)** Adult and **(C)** Aged WT (orange) and *Gata2^+/−^* (green) HSCs. Each dot represents the transcriptome of and individual sample. **(D)** Hallmark GSEA of *Myc* targets in E14 **(E)** Adult and **(F)** Aged WT and *Gata2^+/−^* HSCs. (**G**) Commonly upregulated genes in *Myc* Targets V1 GSEA upregulated in all conditions in *Gata2^+/−^* HSCs. **(H)** Proliferation analysis of adult (**I**) and aged WT and *Gata2^fl/+^*HSCs. (J) Proliferation analysis of young and (K) aged WT and *vav-Cre;Gata2^fl/+^*mice 3 days after CFSE staining. **(L)** Quantification of γH2AX signals in aged-WT and aged-*Gata2^+/−^*after transplantation of LK, LSK, HSC, MPP, HPC1 and HPC2 compartments of the BM. **(M-N)** representative image of a WT and *GATA2^+/−^* K562 cell during anaphase of cell cycle with M) indicating a normal proliferation event and N) indicating an aberrant proliferation event indicated by the arrowhead and quantified in **(O)** N = 3 biological replicates, in total 50 anaphases scored for WT and 49 anaphases scored for *GATA2^+/−^*.

*Myc* is a prominent oncogene^43^ and known to upregulate proliferation. The proliferative phenotype of *Gata2^+/−^* HSCs was validated in cell cycle analysis of adult– and aged-*Gata2^+/−^* HSC populations. Both adult and aged *Gata2^+/−^* HSCs showed a significant loss of quiescent G_0_ phase cells and relevant acquisition of cells in the G_1_ phase of cell cycle, resulting in overall loss of quiescence in the HSC compartment (**Figure 5H-I)**.

Next, LSK cells from *vav-cre;Gata2^fl/+^* adult and aged mice were cultured for 72 hours. CFSE assays revealed a delayed cell cycle progression with fewer cell divisions completed within 3 days of culture in adult mice and was more prominent in aged *vav-cre;Gata2^fl/+^* LSK compared to control **(Figure 5J-K).** Given that GATA2 haploinsufficiency resulted in activation of proliferation related gene sets in all our animal models, the delayed cell cycle progression was unexpected and pointed towards a defect in cell cycle execution. Indeed, RNA-seq of LSK cells 72 hours after proliferation, followed by GSEA, revealed a tendency to a deregulation in pathways involved in DNA replication, chromatin organization and sister chromatid segregation **(Suppl. Figure 6A)**. Additionally, while freshly isolated LSK cells showed only minor genotype-dependent differences (**Suppl. Figure 6B)**, this was increased 3 days after proliferation (FDR-adjusted p-value <0.05: 1974 genes) **(Suppl. Figure 6C),** indicating that hematopoietic cells can compensate for GATA2 haploinsufficiency better under steady-state conditions than under proliferative stress.

On a functional level, we observed significantly increased γH2AX signal in HSCs and HPC1 cells of aged *Gata2^+/−^* BM after transplantation compared to WT, pointing towards the increased presence of DNA double strand breaks (**Figure 5L)**. Together with the chromosomal aberrations found in leukemia (**Figure 3H)**, these data indicate the propensity for genomic instability in GATA2 haploinsufficient cells when forced to proliferate *in vivo* or *in vitro*, supporting the hypothesis that accumulated DNA damage drives leukemic progression in these animals.

To further test this, we generated *GATA2^+/−^* K562 cell lines. K562 cells are diploid for chromosome 3q, containing the *GATA2* locus. Defects in cytokinesis were quantified, which is a common mechanism for karyotype changes resulting in cancer^44^ (**Figure 5M-N**). A higher percentage of *GATA2^+/−^* cells showed defects in cytokinesis compared to GATA2 WT-K562 cells **(Figure 5O**) indicating a direct effect of GATA2 haploinsufficiency on chromosomal segregation and a potential mechanism for leukemic transformation in GATA2 deficiency syndrome. Taken together, our findings demonstrate that Gata2-deficient mouse models need proliferative stress to develop hematological malignancies, that HSCs exhibit impaired transcriptional responses to DNA damage, and that GATA2 heterozygous deletion in cell models leads to chromosomal segregation defects during proliferation.

## Discussion

Patients with GATA2 deficiency exhibit variable cytopenias and increased risk of MDS/AML^2,45–48^. Despite the high prevalence of both hematological phenotypes, the connection between them remains unclear. Here, we have characterized two GATA2 haploinsufficient mouse models that recapitulate various patient specific aspects. Beside cytopenias, some mice presented lethal BMF with severe BM aplasia. Cytopenias arose due to forced proliferation and differentiation defects and, as shown for B cells, increased senescence. Our data shows that ageing and limiting dilution transplantation contributed to disease progression. Aging by itself did not result in these cytopenias; however, *Gata2^+/−^* cells were unable to handle forced proliferation effectively, a characteristic that became more evident in aged cells. In contrast to earlier mouse models^49–51^, leukemias emerged in the absence of induced genetic overexpression of oncogenes. However, leukemia arose exclusively secondary to BMF, underscoring a causal link between these two hematological phenotypes.

Unlike in GATA2-deficient patients, typically developing myeloid neoplasia, the mice presented with T cell leukemias. This difference may be attributed to the high propensity of BL/6 mice to develop immature T cell lymphomas/leukemias^52^. Nevertheless, the model allowed the investigation of different mechanisms involved in malignant transformation. HSCs underwent significant stress like ageing and (limiting dilution) transplantation before leukemic transformation arose, whereas patients develop MDS spontaneously. This difference may be due to the fact that mouse cells have longer telomeres than humans^53^, protecting them from DNA damage, requiring a higher level of proliferative stress to become exhausted.

RNA sequencing showed robust upregulation of *Myc* downstream targets in *Gata2* heterozygous mutant animals compared to WT. *Myc* is a well-known oncogene leading to increased proliferation and reducing checkpoint inhibitors, allowing defective cells in cell cycle and progression to cancer^43^. Indeed, leukemic transformation and MDS were accompanied by chromosomal aberrations and somatic mutations, indicating that germline GATA2 deficiency alone was not sufficient for transformation. Enforced *Myc* expression in HSCs was shown earlier to result in self-renewal of HSCs^54^. It is thus conceivable that increased *Myc* signaling is a survival mechanism to overcome BMF leading to genomic instability. In a zebrafish model for GATA2 deficiency (*gata2a^i^*^4^*^/i^*^4^) NPM1 was identified as a possible target of GATA2 and could result in DNA damage^31^, and it was shown that NPM1 binds directly to MYC, regulating hyperproliferation and malignant transformation^55,56^. Furthermore, our *in vitro* model revealed chromosomal segregation defects and GSEA showed increased expression of genes related to DNA damage. This is in line with the high prevalence of monosomy 7 and trisomy 8 in human MDS with GATA2 deficiency^57^. Further research is required to understand the precise connection between MYC activity and GATA2 deficiency, and how MYC/NPM1 activity levels influence the disease phenotype, including BMF and leukemia. One of the remaining questions is why only part of the mice succumbs post transplantation. Possibly, somatic events compensate for the germline GATA2 defect, like epigenetic changes or increased expression from the WT allele sufficient to maintain hematopoiesis throughout life. A mechanism where the WT allele is upregulated is also seen in a zebrafish model for GATA2 deficiency (*Gata2b^+/−^)*^29^, although this resulted in overcompensation of the *Gata2b* dose along lineage differentiation, resulting in a dysplastic hematopoietic system in the zebrafish kidney marrow (BM orthologue). Allele specific expression of *GATA2* has also been reported in *de novo* AML with acquired GATA2 mutations. These, however, are not null-alleles and it is the mutated allele that is often expressed, suggesting a positive selection mechanism for allele specific expression^58^. It is possible that missense mutations in the ZF2 of GATA2, common in GATA2 deficiency patients, confer novel, oncogenic functions that can induce malignant transformation independently of BMF. In all genetic conditions, however, the germline mutation alone seems not to be enough for the development of myeloid neoplasia. Understanding the nature of these somatic events, including epigenetic modifications, and differentiating between adaptive and maladaptive events will be crucial to modify the disease course and prevent MDS and leukemia.

In conclusion, this study has established that GATA2 haploinsufficiency results in increased *Myc* target expression, a poor proliferative stress response and reduced fitness of HSCs, resulting in cytokinesis defects and leukemic transformation.

## Acknowledgments

We thank U. Kern for insightful discussions. N. Kaltenbach for excellent assistance, and N. Krause and her team of the Center for Experimental Models and Transgenic Services (CEMT) for animal care. We are grateful to M. Follo and her team at the Lighthouse Fluorescence Technologies Core Facility, Freiburg, for cell sorting and maintenance of flow cytometers, Elaine Dzierzak for providing us with *vav-cre;Gata2^fl/+^* mice. ME received support from the European Research Council (ERC Starting Grant no. 638145 “ApoptoMDS” to ME), the German Federal Ministry of Education and Research (BMBF), Berlin (“MyPred – Network for young individuals with syndromes predisposing to myeloid malignancies” no. 01GM1911A), the EJP-RD program (RiboEurope consortium). JF-O is supported by the Deutsche Forschungsgemeinchaft (no. GZ:FE2257/1-1) and by the Hans A. Krebs Medical Scientist Program, Faculty of Medicine, Freiburg, Germany. EdP is supported by KWF/Alpe d’Huzes (SK10321) and EHA junior and senior research grants. We acknowledge funding from the Deutsche Forschungsgemeinschaft (DFG) within the CRC1160 (Project ID 256073931-Z02 to MB), CRC/TRR167 (Project ID 259373024-Z01, MB), CRC1453 (Project ID 431984000-S1, M.B.), CRC1479 (Project ID: 441891347-S1 to MB), TRR 359 (Project ID 491676693-Z01, M.B.), FOR 5476 UcarE (Project ID 493802833-P7, M.B.). We also acknowledge funding from the German Federal Ministry of Education and Research (BMBF) within the Medical Informatics Funding Scheme, PM4Onco–FKZ 01ZZ2322A (MB) and EkoEstMed–FKZ 01ZZ2015 (GA).

## Authorship contributions

JFO, CK, JMW, EG, EdP and ME designed, performed and analyzed experiments; IGM, KG and LQM performed and analyzed the histopathology; GA, MB, RM-L and MS analyzed WES/WGS data, JFO, CK, JMW, EG, BY, CW, HdL, MtB, JZ, RH, KG, EB, SP, CM, SB, MW, EdP and ME analyzed and interpreted data; CN, MR and ME provided patient material; JFO, EdP, CK and ME wrote the manuscript and IP revised the manuscript.

## Figure legends

**Supplementary Figure 1: Gating strategy for flow cytometry.** (A) Gating strategy for HSC populations from BM. B) Apoptosis analysis

**Supplementary Figure 2: *Gata2^fl/+^* and *vav-Cre;Gata2^fl/+^*mice do not show differences in blood cell subsets or differentiation.** Number of stem and progenitor cells in adult **(A)** and aged mice **(B)** in *Gata2^fl/+^* and *vav-Cre;Gata2^fl/+^* mice. **(C)** Colony forming assays from WT and *Gata2^f+/−^* HSC cells and replating **(D)**. Mast cells expansion found on methocults from *vav-Cre;Gata2^fl/+^* mice. **(F)** Serial replating of *Gata2^fl/+^*and *vav-Cre;Gata2^fl/+^* LSK cells.

**Supplementary Figure 3: Serial transplantation of WT and *Gata2^f+/−^*/Gata2^fl/+^ and vav-Cre;Gata2^fl/+^ mice. (A)** Gating strategy of the pro-erythroblast and erythroblast compartments in BM of primary recipients of aged WT and *Gata2^+/−^*cells. Quantified in **(B). (C)** BM cellularity of transplanted mice with aged WT and *Gata2^+/−^* cells. **(D)** Representative example of the serial transplantation assay performed with *Gata2^fl/+^* and *vav-Cre;Gata2^fl/+^* mice. Absolute cell numbers of spleen **(D)**, LN **(E)** and BM **(F)** after 4 serial transplantations.

**Supplementary Figure 4: BMF and leukemia occurs in some recipients transplanted with *Gata2^fl/+^* and *vav-Cre;Gata2^fl/+^* LSK cells. (A)** Analysis of the different BM cell subsets from animals transplanted with both adult and aged *vav-Cre;Gata2^fl/+^* LSK cells that succumbed due to BMF and leukemia compared to *Gata2^fl/+^* mice. **(B)** HSC BM composition of animals transplanted with adult *Gata2^fl/+^* and *vav-Cre;Gata2^fl/+^* LSK cells that remain healthy and were sacrificed after 1 year. **(C)** Blood cell subsets of animals transplanted with adult *Gata2^fl/+^* and *vav-Cre;Gata2^fl/+^* LSK cells that remain healthy and were sacrificed after 1 year. **(D)** Recipient’s pathologies after transplantation with *Gata2^fl/+^* and *vav-Cre;Gata2^fl/+^* LSK cells. % of cells in the SubG0 **(E)** or G0, G1, S and G2/M stages **(F)** 42 days after transplantation.

**Supplementary Figure 5: *Gata2^+/−^* HSCs are marked by proliferative transcriptomic signatures. A)** Network analysis using Cytoscape comparing aged-WT and aged-*Gata2^+/−^* HSCs. Red dots show upregulated gene-sets in *Gata2^+/−^*HSCs compared to WT. Blue nodes are downregulated gene sets and red nodes are up-regulated gene sets connected by lines if differentially regulated genes are in common. **(B)** % of mitotic cell stages performed in adult (left) and aged (right) *Gata2^fl/+^* and *vav-Cre; Gata2^fl/+^* mice.

**Supplementary Figure 6: Forced proliferation leads to gene deregulation in *Gata2^fl/+^* and *vav-Cre;Gata2^fl/+^*LSK cells. (A)** Barplots of upregulated genes (Q-value) 3 days after forced LSK proliferation. **(B)** Volcano plots of freshly and forced to proliferate LSK isolated cells. Genes below the significance threshold are shown in red.

