## Supplementary figures and tables for "Hematological phenotypes in GATA2 deficiency syndrome arise from secondary injuries and maladaptation to proliferation"

### Supplementary Figure 1

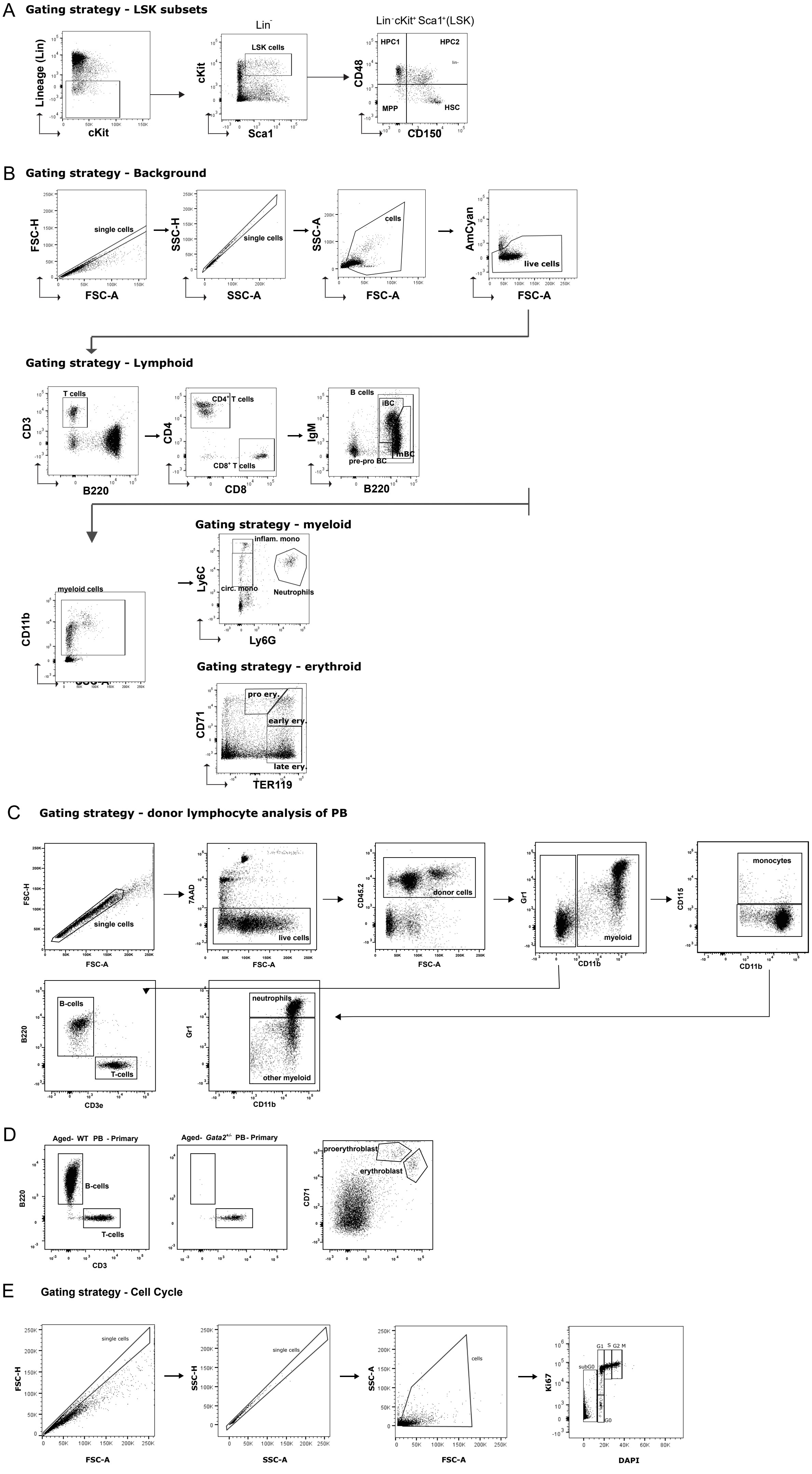

### Supplementary Figure 2

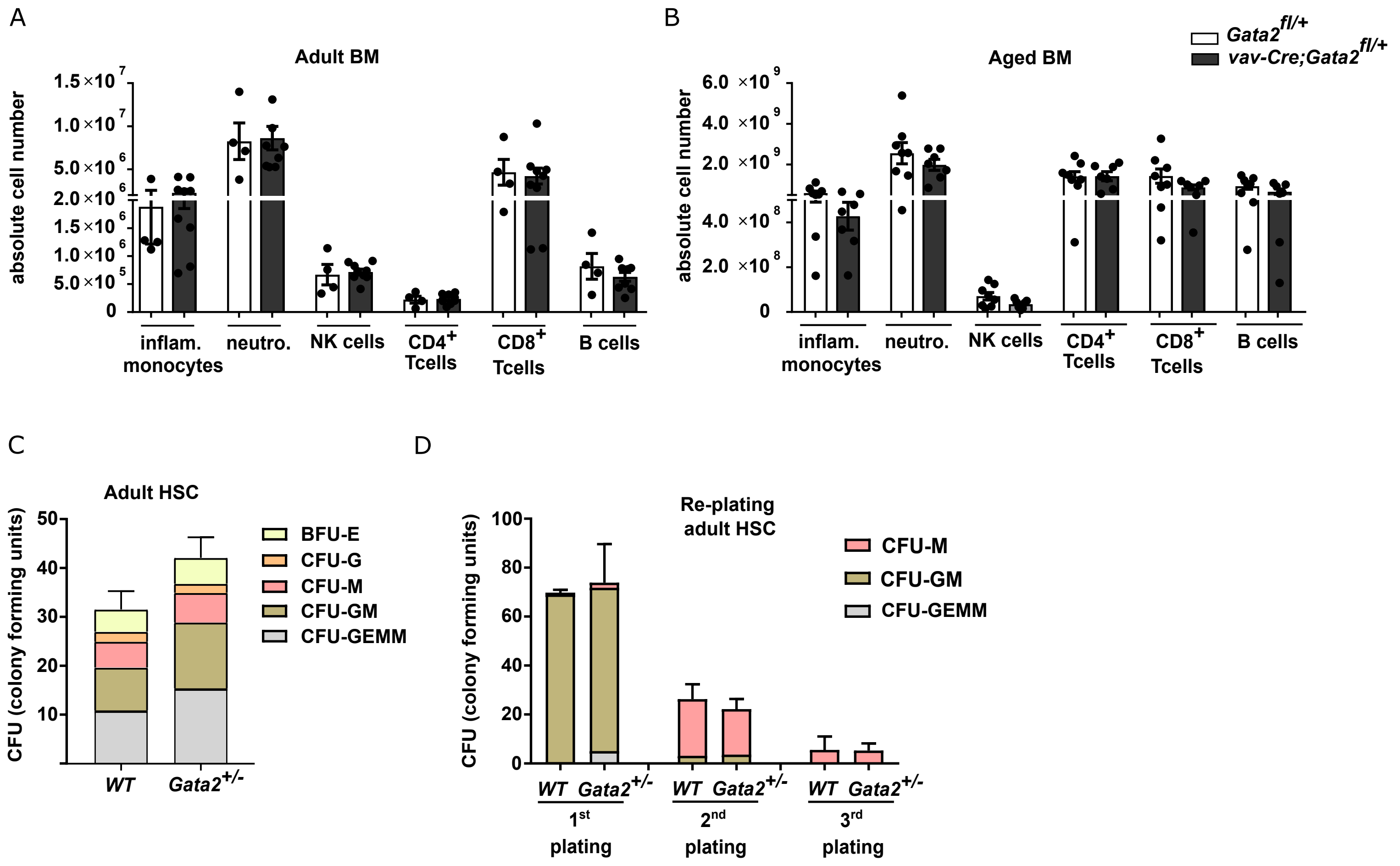

### Supplementary Figure 3

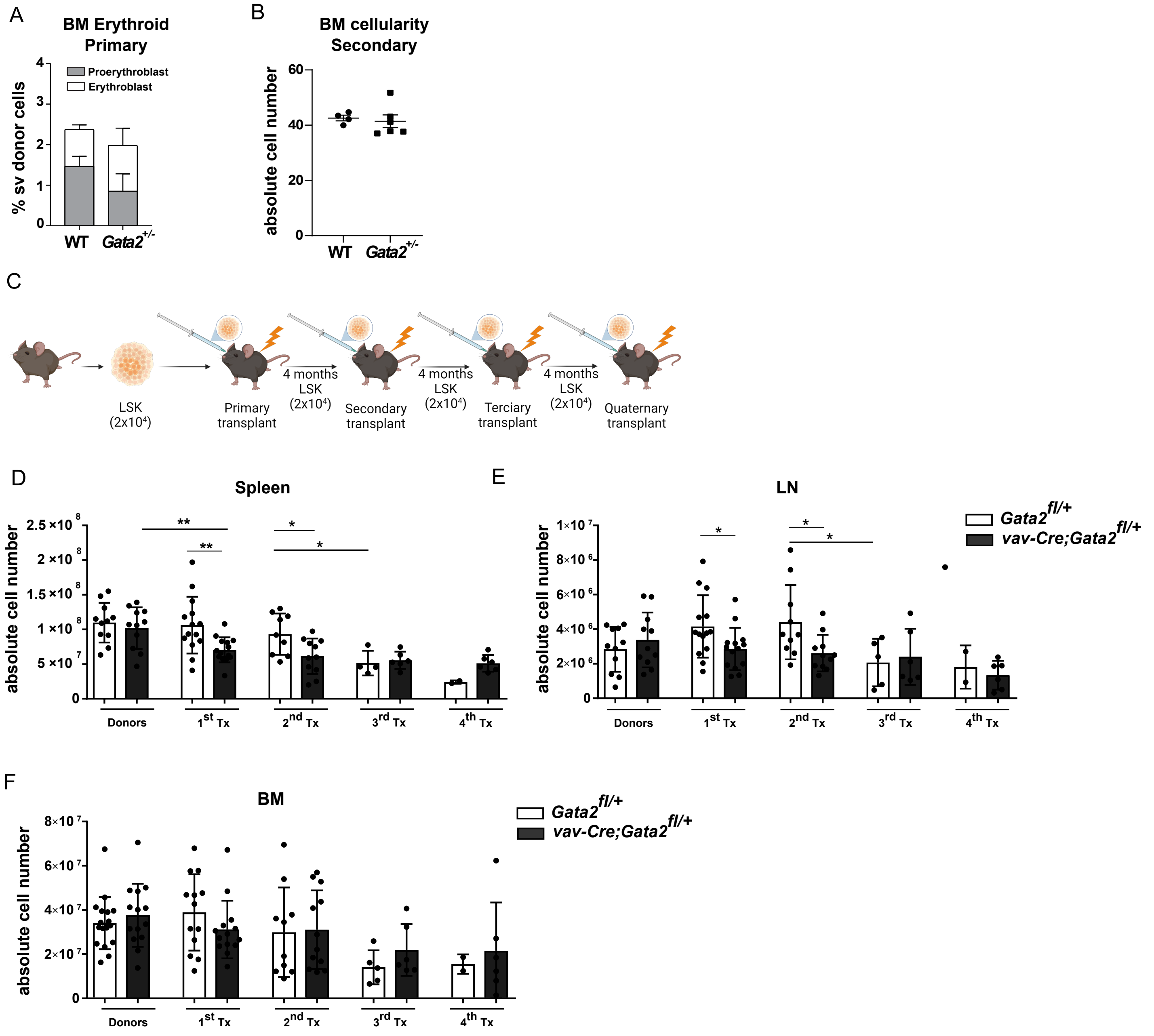

### Supplementary Figure 4

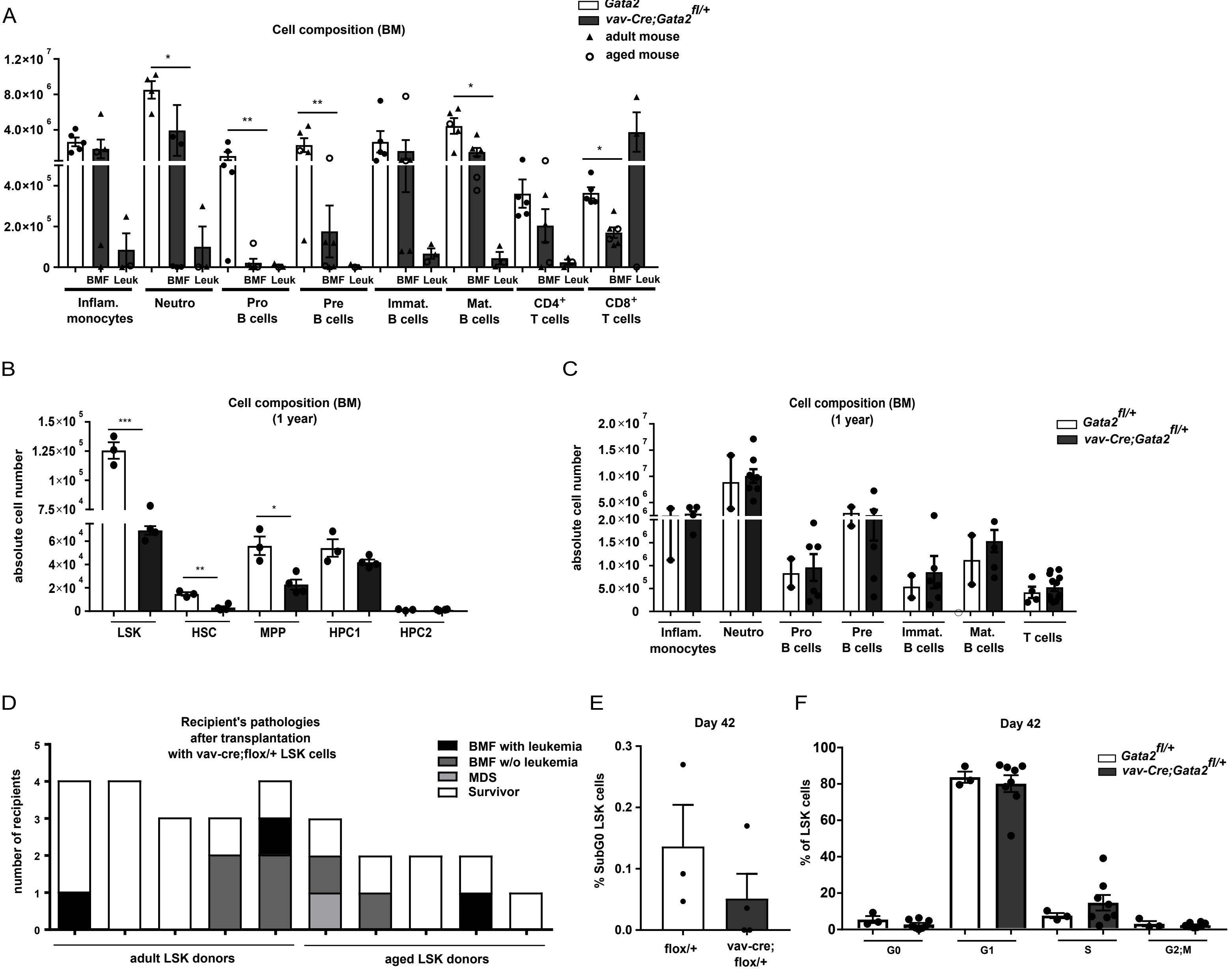

### Supplementary Figure 5

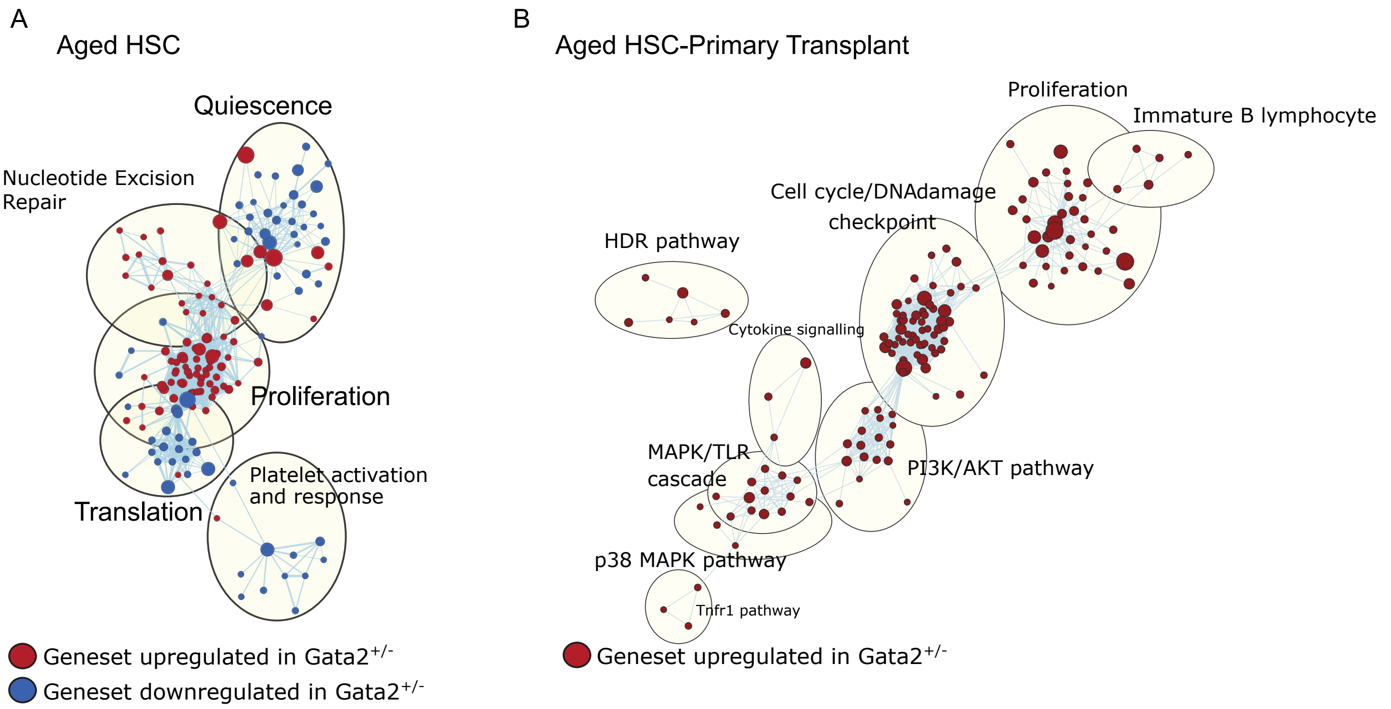

### Supplementary Figure 6

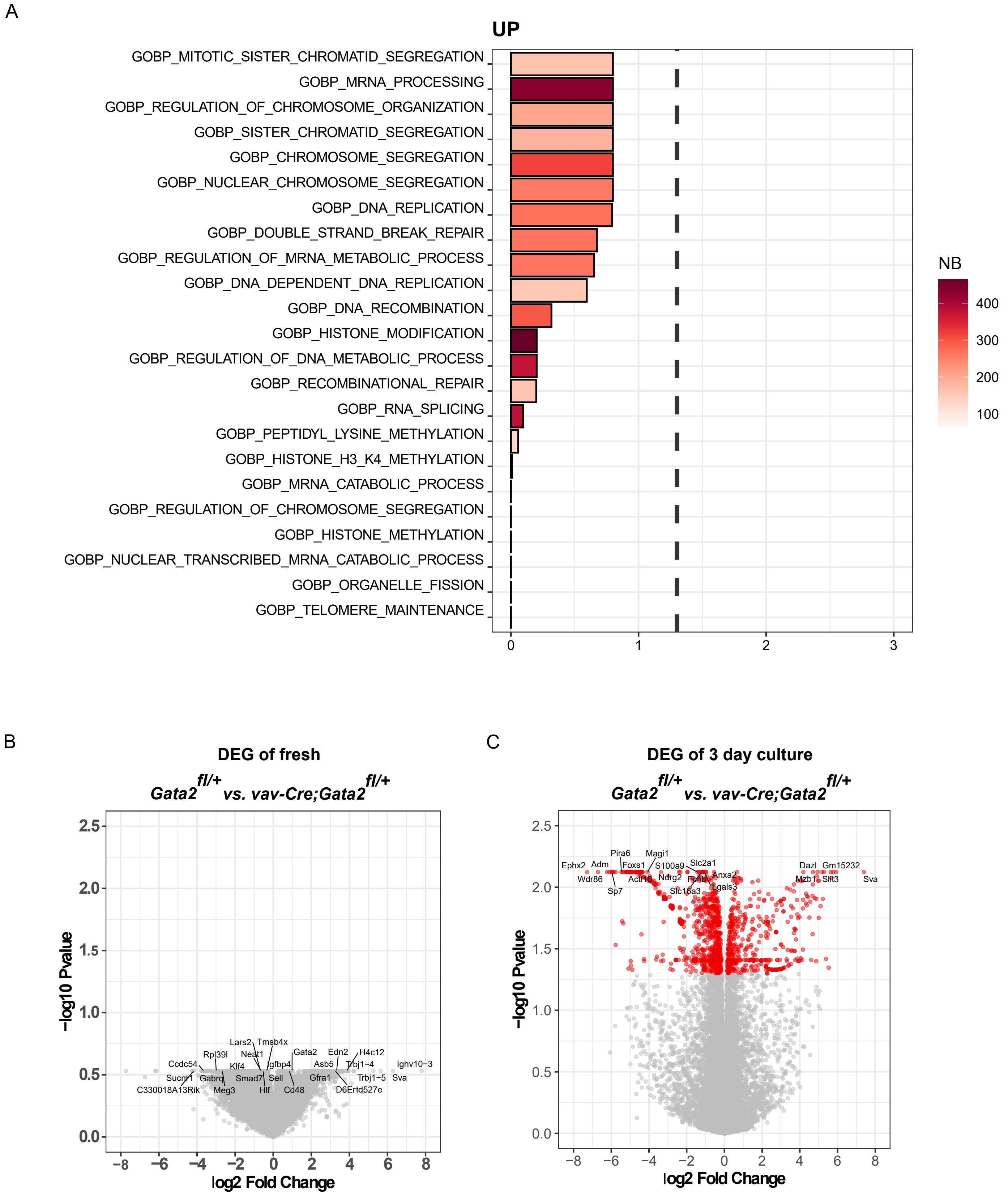

### Supplementary table 1: primers for sequencing

| | |
|---|---|
| **Gata2$^{+/-}$** | |
| **WT forward** | GGA ACG CCA ACG GGG AC |
| **reverse** | GCT GGA CAT CTT CCG ATT CCG GGT |
| **Mutated forward** | GAT CTC CTG TCA TCT CAC CTT GCT |
| **Vav-Cre-Gata2** | |
| **forward** | TCC GTG GGA CCT GTT TCC TTA C |
| **reverse** | GCC TGC GTC CTC CAA CAC CTC TAA |

### Supplementary table 2: antibodies

#### Flow cytometry and sorting antibodies

| Specificity | Clone | Labeling | Manufacturer |
|---|---|---|---|
| CD16/32 | 93 | unlabeled | BioLegend |
| B220 | RA3-6B2 | PE/Cy7 | BioLegend |
| IgM | RMM-1 | APC | BioLegend |
| CD43 | S11 | FITC | BioLegend |
| TCRβ | H57-597 | PE | BioLegend |
| CD4 | RM4-5 | biotin | BioLegend |
| CD8 | 53-6.7 | PerCP | BioLegend |
| CD11b | M1/70 | AF®488 | BioLegend |
| Ly6C | HK1.4 | PE/Cy7 | BioLegend |
| Ly6G | 1A8 | BV421™ | BioLegend |
| Ter-119 | TER-119 | PE | BioLegend |
| CD25 | 3C7 | APC | BioLegend |
| CD44 | IM7 | FITC | BioLegend |
| Sca1 | D7 | APC | BioLegend |
| cKit | 2B8 | PE/Cy7 | BioLegend |
| CD48 | HM48-1 | Pacific Blue™ | BioLegend |
| CD150 | PerCP/Cy5.5 | TC15-12F12-2 | BioLegend |
| CD71 | R17217 | APC | eBioscience |
| CD45.1 | A20 | FITC<br>Pacific Blue<br>biotin | BioLegend |
| CD45.2 | 104 | APC-eFluor® 780 | Invitrogen |
| Ki-67 | SolA15 | FITC | eBioscience |
| Streptavidin | | BV711™ | BioLegend |
| pH2AX(Ser139) | CR55T33 | PE | eBioscience |

### Lineage marker antibodies

| Specificity | Clone | Labeling | Manufacturer |
|---|---|---|---|
| CD3ε | 145-2C11 | biotin | BioLegend |
| CD11b | M1/70 | biotin | BioLegend |
| B220 | RA3-6B2 | biotin | BioLegend |
| Ter-119 | TER-119 | biotin | BioLegend |
| Gr-1 | RB6-8C5 | biotin | BioLegend |
| Nk-1.1 | PK136 | biotin | BioLegend |

### Immunohistochemistry antibodies

| Specificity | Clone | Manufacturer |
|---|---|---|
| CD3 | SP7 | DCS Innovative Diagnostik-Systeme GmbH u. Co. |
| MPO Ab-1 | - | Lab Vision UK |
| Ki67 | - | Abcam plc |
| Ter-119 | - | DakoCytomation |
| Gr-1Ba | - | EMFRET Analytics |
